# Biodiversity genomics of small metazoans: high quality *de novo* genomes from single specimens of field-collected and ethanol-preserved springtails

**DOI:** 10.1101/2020.08.10.244541

**Authors:** Clément Schneider, Christian Woehle, Carola Greve, Cyrille A. D’Haese, Magnus Wolf, Axel Janke, Miklós Bálint, Bruno Hüttel

## Abstract

Genome sequencing of all known eukaryotes on Earth promises unprecedented advances in evolutionary sciences, ecology, systematics and in biodiversity-related applied fields such as environmental management and natural product research. Advances in DNA sequencing technologies make genome sequencing feasible for many non-genetic model species. However, genome sequencing today relies on large quantities of high quality, high molecular weight (HMW) DNA which is mostly obtained from fresh tissues. This is problematic for biodiversity genomics of Metazoa as most species are small and yield minute amounts of DNA. Furthermore, briging living specimens to the lab bench not realistic for the majority of species.

Here we overcome those difficulties by sequencing two species of springtails (Collembola) from single specimens preserved in ethanol. We used a newly developed, genome-wide amplification-based protocol to generate PacBio libraries for HiFi long-read sequencing.

The assembled genomes were highly continuous. They can be considered complete as we recovered over 95% of BUSCOs. Genome-wide amplification does not seem to bias genome recovery. Presence of almost complete copies of the mitochondrial genome in the nuclear genome were pitfalls for automatic assemblers. The genomes fit well into an existing phylogeny of springtails. A neotype is designated for one of the species, blending genome sequencing and creation of taxonomic references.

Our study shows that it is possible to obtain high quality genomes from small, field-preserved sub-millimeter metazoans, thus making their vast diversity accessible to the fields of genomics.

## INTRODUCTION

Biodiversity genomics uses genome-scale data to study the molecular basis of biodiversity. New results already revolutionize life- and environmental sciences by addressing scientific questions about evolution, phylogeny, ecology, etc., assisting the management of biodiversity-related resources, typically through monitoring some aspects of biodiversity (community composition, intraspecific diversity, gene flow, etc.), and through bioprospecting for new resources, e.g. bioactive compounds. Large-scale biodiversity genomics is increasingly possible today, thanks to rapid advances in DNA sequencing.

Two years ago, the Earth BioGenome Project announced plans to sequence all known ~1.5 M eukaryotic species (Lewin et al., 2018). Regarding the taxonomic distribution of these eukaryotes Robert M. May wrote in 1986: “to a good approximation, all species are insects” (May, 1986). This points to an important issue in biodiversity genomics: most known eukaryotic biodiversity belongs to small species. Considering only metazoans, several million species remain to be described (Stork, 2018), and we have good reasons to expect that the absolute majority of these are also in the below one mm size range (Stork, McBroom & Hamilton 2015). The ability to genome-sequence small metazoans is thus necessary for biodiversity genomics.

However, minute animals pose some real challenges to genome sequencing, starting with specimens collection. For example, soil microarthropods are generally field-trapped (e.g. with pitfall trap), extracted from their habitat—e.g. with a Berlese funnel (Berlese, 1904) or subsequently improved systems such as Macfadyen (1961). Capture on sight (e.g. with a mouth aspirator) is tedious and used to a lesser extent. Regardless of the method, members of a sampled community are collectively gathered in a collection tube. Keeping small animals extracted from their environment alive is difficult as they are fragile, sensitive to heat, desiccation, and they might prey on or defend from each other. In most instances, there are incompressible delays between specimens collection and DNA extraction: trapping time, field trip schedule and travel time, collected material sorting, identification requiring specimens preparation for microscopy and morphological analysis itself. Unexpected delays might also occur (need to repeat a failed lab work for example). Therefore the specimens must generally be preserved on the spot to avoid their loss during processing time.

It is often the case that only a few specimens of a given species are brought back from a field trip. Also, the possibility of high level of genetic diversity within a population or of sympatric occurrence of divergent genetic lineages within an asexual morphospecies (e.g. Montero-Pau, Ramos-Rodríguez, Serra, & Gómez 2011; Schneider & D’Haese 2013) increases the complexity of obtaining an accurate haploid genome assembly from pooled specimens. Clonal amplification or in-breeding could overcome this issue, but setting up new cultures for each different species is not realistic. Therefore, except for a handful of very common or laboratory-cultured species (such as the springtail *Folsomia Candida*, a soil-dwelling model species sequenced by Faddeeva-Vakhrusheva et al., 2017), working from a large amount of fresh tissue is simply not an option for biodiversity genomics of small metazoans. Genome sequencing from a single, field-preserved specimen alleviates those obstacles.

Several library preparation approaches were developed for short-read sequencing on Illumina platforms from minute DNA quantities (Carøe et al., 2017; Meyer & Kircher, 2010). Short reads alone are not sufficient to produce low-fragmented genome assemblies, so Illumina-sequenced genomes are not sufficiently high quality for many applications (Jarvis, 2016). Today the technical workhorses for highly continuous genomes are long-read DNA sequencers, produced by two companies: Pacific Biosciences (PacBio), and Oxford Nanopore Technologies (ONT). Long-read sequencing requires high molecular weight DNA, available in high quantities. The first precondition is obviously not negotiable, but much effort is directed to decrease the amount of necessary DNA. For example, PacBio recently released a “Low input library preparation” protocol, with a recommended requirement of 150 ng DNA (https://www.pacb.com/wp-content/uploads/Procedure-Checklist-Preparing-HiFi-Libraries-from-Low-DNA-Input-Using-SMRTbell-Express-Template-Prep-Kit-2.0.pdf), down to 50 ng DNA

(Kingan et al. 2019a). This allows amplification free genome-sequencing of small metazoans in the size range of single mosquitoes (Kingan et al., 2019b, 2019c). But as most metazoans are smaller than a single mosquito, this 50--150 ng DNA requirement will generally prove to be already too high.

Whole genome amplification (WGA) is frequently used to amplify the genomes of unicellular unculturable organisms (Binga, Lasken, & Neufeld 2008; Woyke, Doud & Schulz 2017), including unicellular eukaryotes (Fischer et al., 2009). Although WGA was proposed for small metazoans long ago (Gorrochotegui-Escalante & Iv, 2003), we are aware of only a few applications to date. For example, several springtail genomes were Illumina-sequenced after WGA (Sun, Ding, Orr & Zhang, (2020). Here we present results from a new, ultra-low input (UL) library protocol from Pacific Biosciences, which allows as little as 5 ng gDNA input to be used for long read sequencing. We successfully created UL sequencing libraries from two springtail species: *Desoria tigrina* Nicolet, 1842 (2 mm; Fig. 1A) and *Sminthurides aquaticus* (Bourlet, 1841) (1 mm; Fig. 1B–E), and sequenced these on PacBio Sequel II platforms. WGA is not without problems: amplification bias and chimera formation might reduce the uniformity in sequencing coverage and genome completeness (Pinard et al., 2006; de Bourcy et al., 2014). We evaluated these issues by sequencing a pool of several specimens of *S. aquaticus* on PacBio Sequel II, after a non-genome-wide amplification-based low input library preparation (Kingan et al., 2019b).

**Figure 1.**
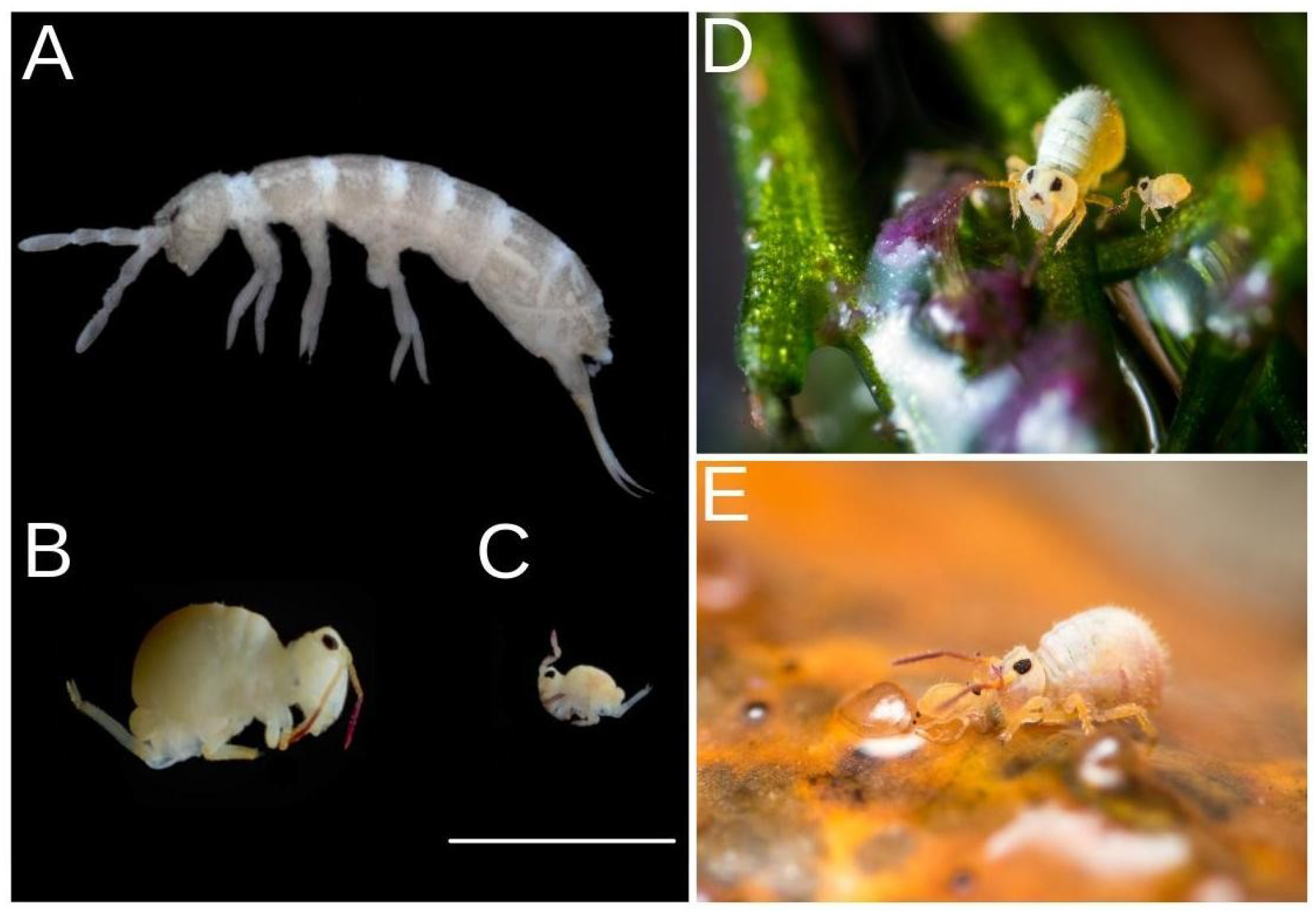
(A) *Desoria tigrina. Sminthurides aquaticus* (B) female, (C) male, (D) male and female on wet plant, (E) courtship on a floating dead twig: the male uses its clasping antennae to grab the antennae of the much bigger female. (A–C) Specimens preserved in 96% ethanol, scale bar = 1 mm.

*D. tigrina* (Entomobyomorpha, Isotomidae) is an epedaphous species: upper layer of soil and litter dweller mostly found in anthropized environments (Potapov, 2001). It can be very abundant in vegetal compost, is also found in crop fields (Gruss & Twardowski, 2016), and can occur in caves as a troglophile (Dányi 2011). In western europe it remains active in winter. It has been reported from Macquarie Island (sub-antarctic island), where it is believed to be alien but not found more tolerant to low or high temperatures than the indigenous species (Phillips et al., 2020). *S. aquaticus* (Symphypleona, Sminthurididae) is an epigeous, hygrophilous species widely spread in the Holarctic region (Bretfeld 1999). The species is common in fresh waters. It gathers on solid ground created by emerging plants, wood and rocks and can efficiently walk and jump on water surfaces thanks to its elongated claws and strong furca (jump appendage) with tip (mucro) shaped as a paddle. The species is also remarkable by its strong sexual dimorphism: the male is significantly smaller than the female and its modified antennae into a prehensile organ allows it to clasp the female antenna in a courtship dance preceding external fecondation (Fig. 1 B–E). It is a relatively well studied species according to Collembola standards. Its body mass is estimated around 0.05 and 0.14 mg (Eisenberg, 1989). A peculiar postzygotic of sex determination through aberrant spermatogenesis was described by Dallai, Fanciulli & Frati (2004).

## MATERIALS AND METHODS

### Specimens collection and preparation

*D. tigrina* was collected in a compost bin, from a garden in Germany, Hesse, Sulzbach am Taunus; X=8.5213°, Y=50.1393°, 14.xii.2019. Specimens were extracted from the compost with a Berlese funnel, and falled directly in 96% ethanol. DNA extraction was performed within ~72h.*S. aquaticus* was collected from a pond in a public garden: France, Île-de-France, Paris, Jardin Naturel Pierre-Emmanuel, X=2.3999°, Y=48.8589°, 9.i.2015, 27.x.2019, 29.vii.2020. Specimens were eye-caught using a small net, and fixed in 96% ethanol. The specimens from the 2019 collection were preserved in 96% ethanol at −20°C until DNA extraction. Specimens used for morphological identification were cleared in lactic acid. Specimens were further bleached in a solution of potassium hydroxide (KOH). Specimens were mounted in permanent slides using Marc-André II mounting medium. Observations were made using a Leitz Wetzlar Diaplan with phase contrast, at 400-1000x magnification.

### PacBio ultra-low input (UL) library preparation

Extraction was performed from a single specimen each time. Specimen was washed in 1 x PBS (Sigma) to remove as much as possible residual EtOH. EtOH was first replaced with PBS, then the solution was replaced four times with clean PBS. Specimen was crushed with one-way pistils (Sigma) then DNA was extracted according to the Qiagen MagAttract kit (Hilden, Germany). Altogether, we performed eight extractions from *S. aquaticus* and four extractions from *D. tigrina* specimens. Extracted DNA was quantified with the Quantus dsDNA system (Promega) and quality was assessed by FEMTOpulse (Agilent). UL input libraries were prepared using an early access kit kindly provided by Pacific Biosciences. Briefly, gDNA was fragmented with g-Tubes (Covaris) and the resulting fragment size was again inspected by FEMTOpulse (Agilent). Next, single-stranded overhangs were removed enzymatically, followed by a DNA damage repair, end repair and an A-tailing step. A double-stranded DNA adapter with a T-overhang was ligated for 1 h at 20°C and the resulting products were bead purified (ProNex, Promega), eluted and then split into two identical aliquots. DNA fragments with adapters were amplified by two different PCR reactions (reaction 1: 98°C for 45 s, 14 cycles: 98°C for 10s, 62°C for 15s, 72°C for 7 min, final elongation 72°C 5 min; reaction 2: 98°C for 30 s, 14 cycles: 98°C for 10s, 60°C for 15s, 68°C for 10 min, final elongation 68°C 5 min). PCR reactions were again bead purified, eluted in EB and library fragments were again quantity (Quantus, Promega) and quality assessed (FEMTOpulse, Agilent). PCR fragments from both reactions were pooled in equal amounts to achieve a total of 500 ng input. Next, libraries were prepared as outlined in the low input protocol (https://www.pacb.com/wp-content/uploads/Procedure-Checklist-Preparing-HiFi-Libraries-from-Low-DNA-Input-Using-SMRTbell-Express-Template-Prep-Kit-2.0.pdf). Libraries were then annealed to a sequencing primer (V4), bound to Sequel II DNA polymerase 2.0 with Binding kit 2.0 and sequenced in a Sequel II 8M SMRT cell for 30 h.

### PacBio low input (L) library preparation

12 specimens of *S. aquaticus* were pooled in 3 groups of four specimens. Genomic DNA was extracted from each pool using the 10X Genomics^®^ (Pleasanton, CA) “Salting Out Method for DNA Extraction from Cells” (CG000116 Rev A, adapted from Miller, Dykes & Polesky 1988) protocol with following modifications: specimens were ground in the lysis buffer with a pestle before incubation, and the completed with a second wash by repeating DNA precipitation. The extract purity was evaluated with the NanoDrop spectrophotometer (Thermo Fisher Scientific, Waltham, MA). DNA was resuspended in 30 μL of TE buffer, and 4 μL was used for concentration measurement using the Qubit Fluorometer and Qubit dsDNA HS reagents Assay kit (Thermo Fisher Scientific, Waltham, MA). We finally combined the three extracts to reach DNA quantities sufficient for library preparation. Fragment size distribution was checked on an Agilent TapeStation (Santa Clara, CA). Combining all extraction pools resulted in a total amount of 195 ng HMW DNA. SMRTbell library was constructed following the instructions of the SMRTbell Express Prep kit v2.0 With Low DNA Input Protocol (Pacific Biosciences, Menlo Park, CA). Total input DNA was approximately 170 ng. Ligation with T-overhang SMRTbell adapters was performed at 20°C overnight. Following ligation, the SMRTbell library was purified with an AMPure PB bead clean up step with 0.45X volume of AMPure PB beads. Subsequently a size-selection step with AMPure PB Beads was performed to remove short SMRTbell templates < 3kb. For this purpose the AMPure PB beads stock solution was diluted with an elution buffer (40% volume/volume) and then added to the DNA sample with 2.2X volume. The size and concentration of the final library were assessed using the TapeStation (Agilent Technologies) and the Qubit Fluorometer and Qubit dsDNA HS reagents Assay kit, respectively. Sequencing primer v4 and Sequel^®^ II Polymerase 2.0 were annealed and bound, respectively, to the SMRTbell library. Due to low sample concentration, the library was loaded at an on-plate concentration of only 14 pM using diffusion loading. SMRT sequencing was performed in CLR mode on the Sequel System II with Sequel II Sequencing Kit 2.0, 30 hour movie time with NO pre-extension and Software SMRTLINK 8.0. A total of 1 SMRT Cells was run. The sequencing was performed at the Radboud University Medical Center, Nijmegen (NL).

### UL sequencing data assembly

Generation of circular consensus sequences (CCS) and adapter trimming was done in PacBio SMRTLink 8 with default parameters followed by deduplication of reads via pbmarkdup (Version 0.2.0, https://github.com/PacificBiosciences/pbmarkdup) as suggested by PacBio for the UL protocol. Further HiFi reads of *D. tigrina* were discarded, if they were found to contain complete UL PCR Adapter sequences or the reverse-complement. Genome properties estimation relying on kmer statistics prior to assembly was possible due to the low error rates of HiFi reads. K-Mer were counted and aggregated using jellyfish 2.2.10 (Marçais & Kingsford, 2011) (‘jellyfish count -C -m 21 -t 20 -s 1000000000 -o jelly_k21.jf CCS.fasta’ & ‘jellyfish histo -t 10 jelly_k21.jf > kmer.histo’). GenoScope 1.0 (Vurture et al., 2017) was used to estimate genome length, level of duplication and heterozygosity through the web application (http://qb.cshl.edu/genomescope/), using the median CCS reads length in parameter. Assemblies were produced via FALCON (Version falcon-kit 1.8.0, parameters adapted: genome_size = 200,000,000, -e.98 (ovlp_daligner_option), --min-idt 98 (overlap_filtering_setting) and polished using racon (Version 1.4.10, parameter: ‘-u’, https://github.com/isovic/racon) in combination with samtools (Version 1.9, parameters: ‘-F 1796 -q 20’; http://www.htslib.org/) and the PacBio wrapper of minimap2 (Version 1.1.0, parameters: ‘--preset CCS --sort’, Li 2018 https://pubmed.ncbi.nlm.nih.gov/29750242/) following PacBio guidelines (https://github.com/PacificBiosciences/pbbioconda/wiki/Assembling-HiFi-data:-FALCON-Unzip3). Further removal of remaining haplotigs was performed using purge_dups (Version 1.0.1, Guan et al., 2020, https://github.com/dfguan/purge_dups) for *D. tigrina* and, with additional removal of artefact contigs, via purge_haplotigs (Version 1.1.0, Roach, Schmidt & Borneman 2018, https://bitbucket.org/mroachawri/purge_haplotigs/src/master/) for *S. aquaticus*. Here we followed the pipelines described at the webpages of the corresponding tool (See above, using minimap2 version 2.17 with parameter ‘-x asm20’ instead of ‘ -x map-pb’). Residual adapter contamination was observed in both dataset. Reads were screened for adapter sequences, and removed. Ncbi-blastn was used to gather the CCS containing exclusively mitochondrial sequences and use them to assemble the mitochondrial genomes using Geneious (2020.1.2). Circularity was validated manually, and nucleotide bases were called with a 75% threshold consensus. The mitochondrial genomes were annotated with MITOS2 web server (Bernt et al., 2013). CDS were checked and corrected using Geneious, to ensure that the presence of uncommon start codon and incomplete stop codons did not mislead the automatic annotation algorithms. Boundaries of the rDNAs were slightly adjusted to make them contiguous with the tRNA(val) gene.

To identify insertions of the mitochondrial genome in the nuclear genome (NUMTs) we used ncbi-blastn with a 2x duplicated sequence of the mitochondrial genome as a query (to handle circularity). We recognized the presence of almost complete copies of the mitochondrial genome in the two species, with length equal or superior to the mean length of the CCS (Figure 3). We investigated the quality of the UL assemblies in those locations using IGV (Robinson et al. 2011) and we applied the following procedure to NUMTs with length > 10 kb: at both extremities of the NUMT, a ~1000 bp sequence with a 500 bp overlap on the contiguous nuclear sequence was selected. All CCS aligning with those sequences were gathered (blastn) and reassembled using Geneious. Matching extremities were bridged by visual recognition of the specific DNA sequence of each NUMT, made possible by the high accuracy of the CCS. Whole genome assemblies were manually corrected by splitting, swapping and resoldering contigs where missaemblies were observed. When a given extremity could not be bridged to another, the contigs were left splitted.

### L sequencing data assembly

The CLRs were assembled alternatively with Flye v2.7.1 (Lin et al., 2016; Kolmogorov, Yuan, Lin & Pevzner, 2019) using default parameters including the polishing step, and FALCON (Chin et al., 2016; version falcon-kit 1.8.1). FALCON was run with adapted parameters: genome_size = 200,000,000; length_cutoff; = 3000; pa_daligner_option = -e0.76 −l1200 -k18 -h70 -w8 -s100; pa_HPCdaligner_option = -v -B128 -M24; pa_HPCTANmask_option = -k18 -h480 -w8 -e.8 -s100; pa_HPCREPmask_option = -k18 -h480 -w8 -e.8 -s100; pa_DBsplit_option = -x500 -s400; falcon_sense_option = --output-multi --min-idt 0.70 --min-cov 3 --max-n-read 400 --n-core 24; length_cutoff_pr = 5000; ovlp_daligner_option = -k24 -h1024 -e.95 -l1800 -s100; ovlp_HPCdaligner_option = -v -B128 -M24; ovlp_DBsplit_option = -s400; overlap_filtering_setting = --max-diff 100 --max-cov 150 --min-cov 3 --n-core 24. The FALCON assembly was polished once with Racon (version 1.4.3, parameter: ‘-u’, https://github.com/isovic/racon) using an alignment produced with minimap2 (version 2.17, parameters: with -H -x map-pb option) and Racon default parameters.

### Contamination control

We checked the UL assemblies for contamination using a set of tools. We use Sourmash (Pierce, Irber, Reiter, Brooks & Brown, 2019) to create minHash signatures of each contig and compare them to pre-made search databases for taxonomic assignment of microbial genomes (bacterial, viral and fungal, Genbank LCA Database 2017.11.07, https://sourmash.readthedocs.io/en/latest/databases.html), using k-mer sizes of 21, 31, 51. We also aligned contigs against the NCBI nr protein database (downloaded on February 7, 2020) with DIAMOND, using long read settings (Buchfink, Xie & Huson, 2015). We evaluated DIAMOND hits with MEGAN 6.18.10 (Huson et al., 2016), using its long read algorithm (Huson et al., 2018). We used the following settings for the MEGAN assignment: minScore 500, maxExpected: 0.01, minPercentIdentify 0, topPercent: 5, minSupportPercent: off, percentToCover 51. Finally, we used ncbi-blastn to query the contigs against the NCBI nr nucleotide database (-max_target_seqs 10 -max_hsps 1 -evalue 1e-25). We used the BlobToolKit pipeline (Challis, Richards, Rajan, Cochrane & Blaxter, 2020) to merge and plot the DIAMOND and BLAST taxonomic assignment with the assembly statistics (GC content and base coverage). Contigs explicitly assigned to anything else than metazoan were excluded from downstream analysis.

### Assemblies assessment

All assemblies completeness was assessed using BUSCO v4.0.6 (Seppey et al., 2019), in genome mode and --long option, with the arthropoda_odb10 dataset (Kriventseva et al., 2019). To evaluate assembly quality and statistics, we used backmap (v0.3 Schell et al. 2017) a perl wrapper of minimap2 and QualiMap (Okonechnikov et al., 2016). minimap2 was run with “-H -x map-pb” to map CLR subreads on the L assembly, and “-H -ax asm20” to map CCS on the UL assembly. backmap estimates genome size from mapped nucleotides divided by mode of the coverage distribution.

### Annotation

UL assemblies of both species were annotated with *ab initio* gene prediction. Repetitive regions were masked in the assembly based on RepeatModeler 2.0.1 (Flynn et al., 2020) with the options: ‘-LTRStruct -engine ncbi’ using RepeatMasker (open-4.0.9, options: ‘-xsmall -gff -nolow‘). Protein sequences were predicted via AUGUSTUS (Version 3.3.3, Stanke et al., 2006, option: ‘--softmasking=on‘) based on BUSCO (Version 4.0.6, ‘--long’ option) training results. Functional annotations were obtained by a local installation of eggNOG-mapper (Version v2.0.1, Huerta-Cepas et al., 2019, option: ‘-m diamond‘). If emapper recovered no annotations, we denoted sequences as ‘hypothetical protein’ (for proteins without hits in emapper), or ‘uncharacterized protein’ (for proteins with hits without annotations). Collembola are considered the only known animal group that might be able of antibiotic synthesis through the horizontal transfer of beta-lactam gene clusters from soil fungi (Faddeeva-Vakhrusheva et al., 2017; Suring et al., 2017). We searched the genomes for genes homologous of the isopenicillin N synthase-like gene found in *F. Candida* by querying the nucleotide sequence of the mRNA (NCBI accession XM_022089550.1) using blastn and DIAMOND blastx against the genome assembly (Buchfink, Xie & Huson, 2015).

### Phylogenetic analysis

We downloaded 14 springtails and 1 dipluran genomes assemblies from NCBI (dataset expended from Sun et al. 2020, table 1). We used BUSCO v4.0.6 in short mode and restricted the ortholog search to BUSCO’s in the arthropoda_odb10 dataset. We performed gene prediction with AUGUSTUS v2.5.5, trained with the common fly dataset. We screened the obtained BUSCO sets with a custom script to identify genes shared among the species, allowing for 25% missing data. We aligned single protein sequences with MAFFT v7.450 (Katoh & Standley, 2013), concatenated the alignments with FASconCAT-G v1.04 (Kück & Longo, 2014), and trimmed the final alignment with trimAl v1.2 (Capella-Gutiérrez, Silla-Martínez & Gabaldón 2009). We calculated a maximum likelihood tree with IQtree v1.6.12 (Nguyen, Schmidt, von Haeseler & Minh, 2015) with 1000 bootstrap replications.

**Table 1.**
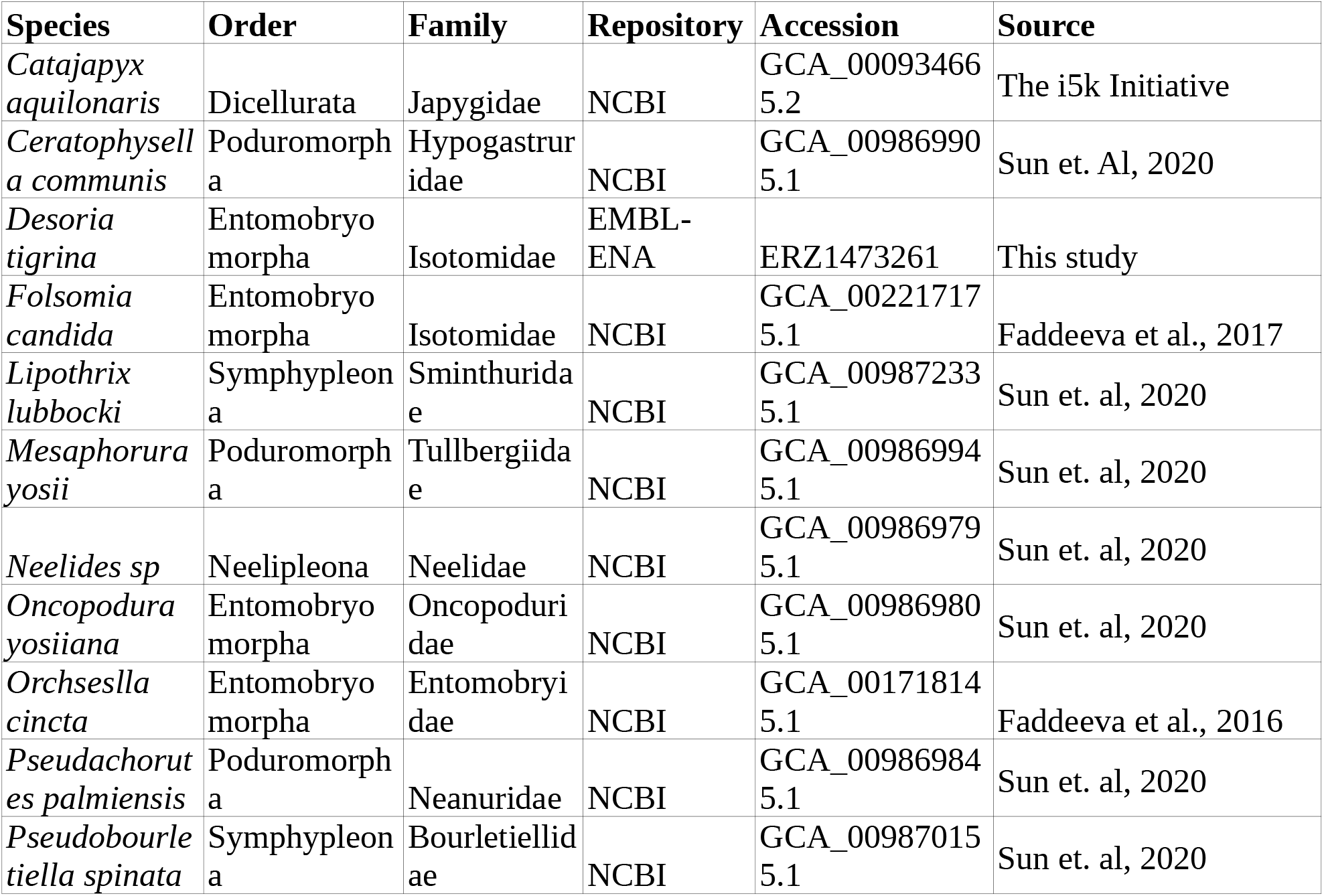
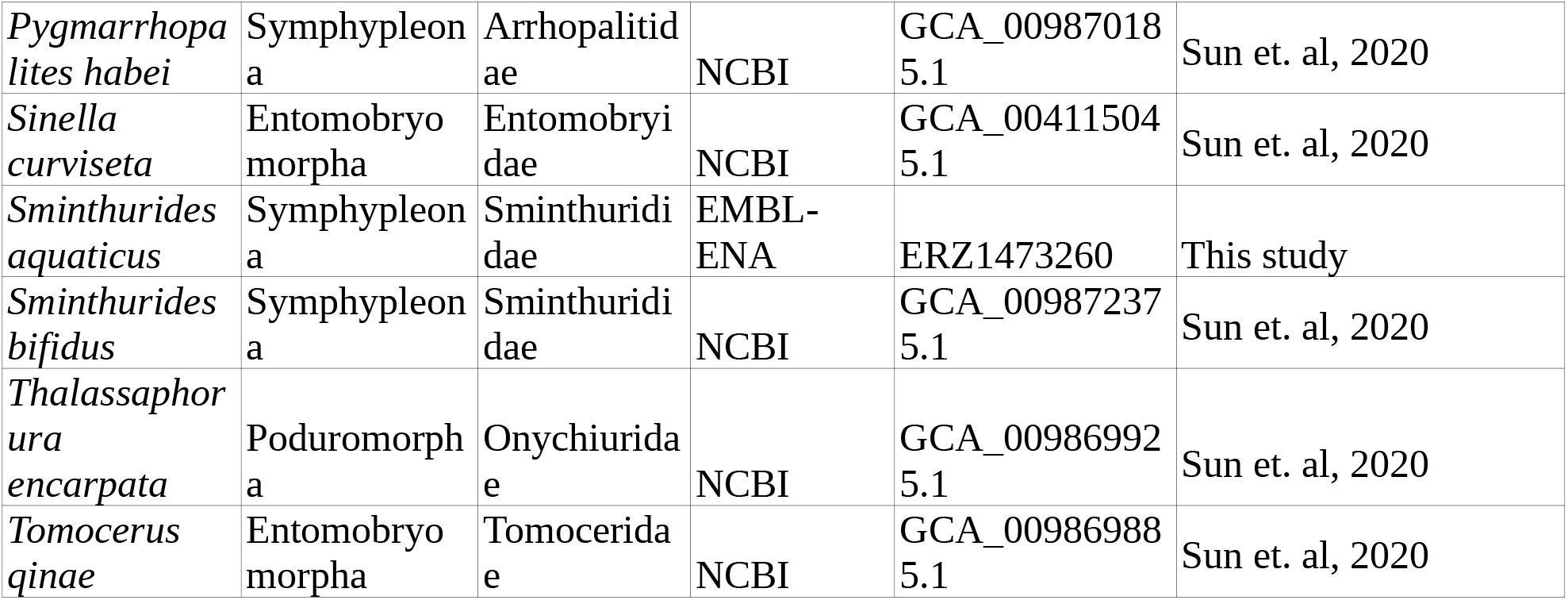
Species included in the phylogenetic analysis (dataset expended from Sun et. al 2020).

## RESULTS

### Species biology and taxonomy

*D. tigrina* was by far the dominant springtail species in the compost bin during the winter season (biomass and number of individuals). The species is still present in summer but specimens are smaller in size and share the habitat with less abundant but larger *Sinella* sp. and *Tomocerus* sp. (not further studied). Morphological observations unambiguously set the collected specimens within the *D. tigrina-group* (Fjellberg 2007). Within this group, outer maxillary palp chaetotaxy was used to distinguish *D. tigrina* from its sibling species *D. grisea* (Fjellberg 2007). Identification was further validated following Potapov (2001). 17 females and 3 males on 12 slides labelled CSCH-1326—1337 will be deposited to the Apterygota collection of the Senckenberg, Görlitz. 6 females, 2 males and 2 juveniles on 4 slides numbered EA013940-43 will be deposited to the Apterygota collection of the National Museum of Natural History, Paris.

*S. aquaticus* was the only springtail species forming a population on the pond (i.e. no accidental fall on water surface). Abundantly found in October 2019, it was observed again in June 2020 in large numbers and courtship behavior undergoing (Fig 1D, E). The specimens identification was unambiguous following Bretfeld (1999), Fjellberg (2007) and Stach (1956) descriptions. One male on a slide numbered CSCH-1338 is designated as the neotype for *S. aquaticus* (see discussion) and will be deposited to the Apterygota collection at the National Museum of Natural History, Paris, along with 3 females and 2 males on five slides (CSCH-1339–1344) and 20 individuals in 96% ethanol (CS.371, leg. C. Schneider). Three males, three females and one juvenile on 5 slides numbered CSCH-1345–1349 will be deposited to the Apterygota collection at Senckenberg, Görlitz.

### Genome statistics estimations from CCS

For *D. tigrina* a total of 21,3 Gb HiFi data (Q>=20) was produced with a mean read length of 12218 bp with a single 8M SMRT cell. The genome haploid length was estimated to be 167.825 Mbp with 1.43 % of heterozygosity and 3 % duplications. For *S. aquaticus* 13,0 Gb HiFi data with a mean length of 12359 bp was generated and the genome displays lower numbers for those three properties, heterozygosity: 0.96 %, genome size: 152.357 Mbp and duplications 0.78%.

### UL assemblies

For *S. aquaticus*, one step of Purge_haplotigs resulted in a normal distribution of the assembly coverage (no visible diploid peak) and a low BUSCO duplication (2.3 %), suggesting good performance in haplotig purging. For *D. tigrina*, haplotigs purging with Purge_haplotigs resulted in 9.8 % duplicated BUSCOs and Purge_dups resulted in 3.6 % duplicated BUSCOs. The Purge_dups results were kept. *D. tigrina* sequencing achieved a higher mean coverage than *S. aquaticus*, (95.87X vs 71.44X) but also a higher coverage standard deviation. *S. aquaticus* resulted in an assembly of higher contiguity both in terms of N50 and number of contigs. BUSCO completeness is high for both species (*D. tigrina:* 96.1 % complete, *S. aquaticus:* 96.3 % complete).

### L assembly

Flye yielded the best result. This assembly is characterized by a much lower contiguity than the ultra-low assembly, and a high BUSCO (95.8%).

Statistics for all assemblies are reported in table 2, and visuals of UL curated assemblies for both species were created using assembly-stats (version 17.02, https://github.com/rjchallis/assembly-stats) and reported in Figure 2.

**Table 2.**
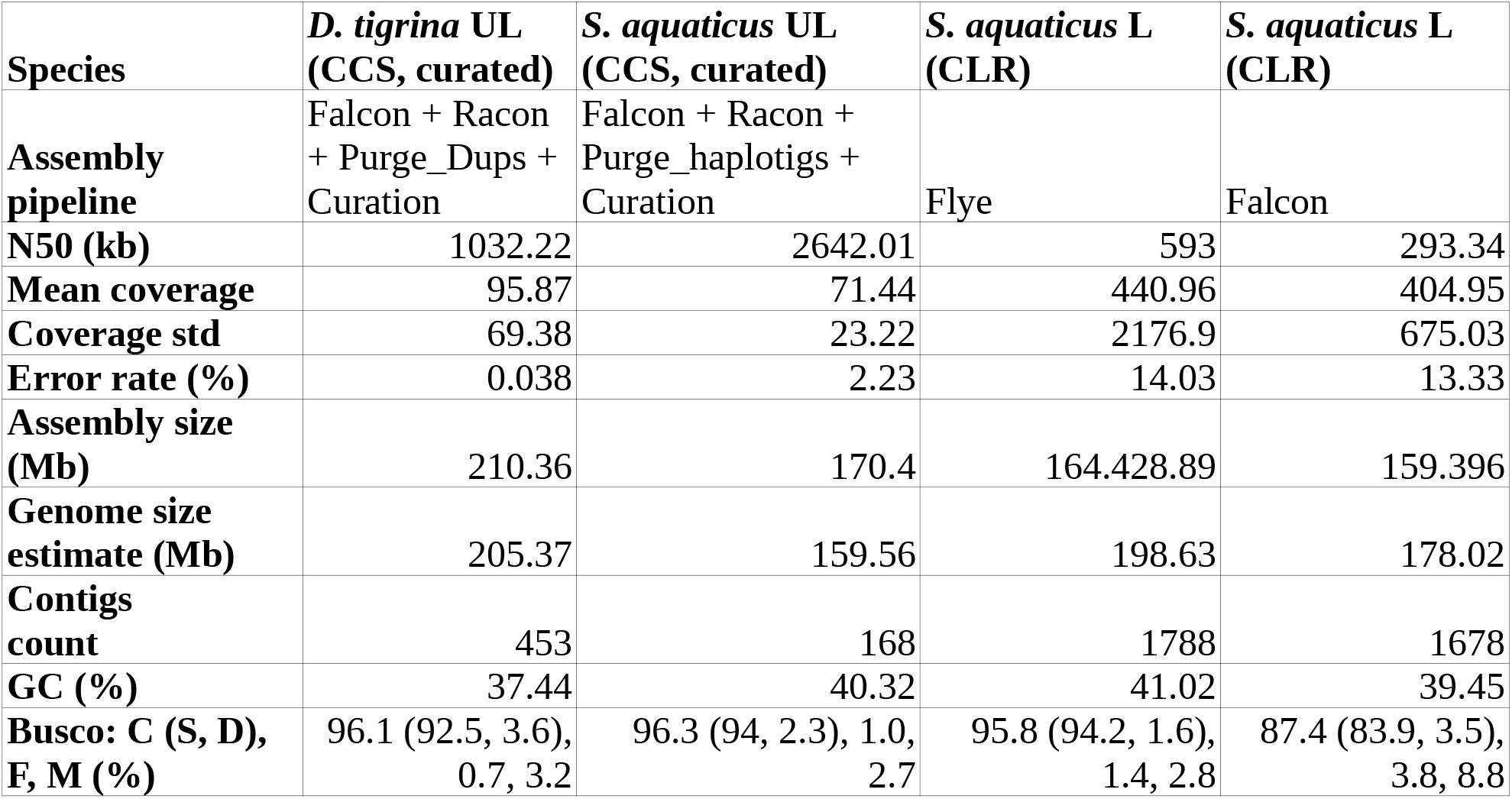
Assemblies statistics. Coverage, error rate and genome size estimation were computed using backmap, Minimap2 and Qualimap. BUSCO scores were computed using BUSCO v4.06 on arthropoda_odb10 with the --long option.

| Species | *D. tigrina* UL<br>(CCS, curated) | *S. aquaticus* UL<br>(CCS, curated) | *S. aquaticus* L<br>(CLR) | *S. aquaticus* L<br>(CLR) |
| --- | --- | --- | --- | --- |
| **Assembly**<br>**pipeline** | Falcon + Racon<br>+ Purge_Dups +<br>Curation | Falcon + Racon +<br>Purge_haplotigs +<br>Curation | Flye | Falcon |
| **N50 (kb)** | 1032.22 | 2642.01 | 593 | 293.34 |
| **Mean coverage** | 95.87 | 71.44 | 440.96 | 404.95 |
| **Coverage std** | 69.38 | 23.22 | 2176.9 | 675.03 |
| **Error rate (%)** | 0.038 | 2.23 | 14.03 | 13.33 |
| **Assembly size**<br>**(Mb)** | 210.36 | 170.4 | 164.428.89 | 159.396 |
| **Genome size**<br>**estimate (Mb)** | 205.37 | 159.56 | 198.63 | 178.02 |
| **Contigs**<br>**count** | 453 | 168 | 1788 | 1678 |
| **GC (%)** | 37.44 | 40.32 | 41.02 | 39.45 |
| **Busco: C (S, D),**<br>**F, M (%)** | 96.1 (92.5, 3.6),<br>0.7, 3.2 | 96.3 (94, 2.3), 1.0,<br>2.7 | 95.8 (94.2, 1.6),<br>1.4, 2.8 | 87.4 (83.9, 3.5),<br>3.8, 8.8 |

**Figure 2.**
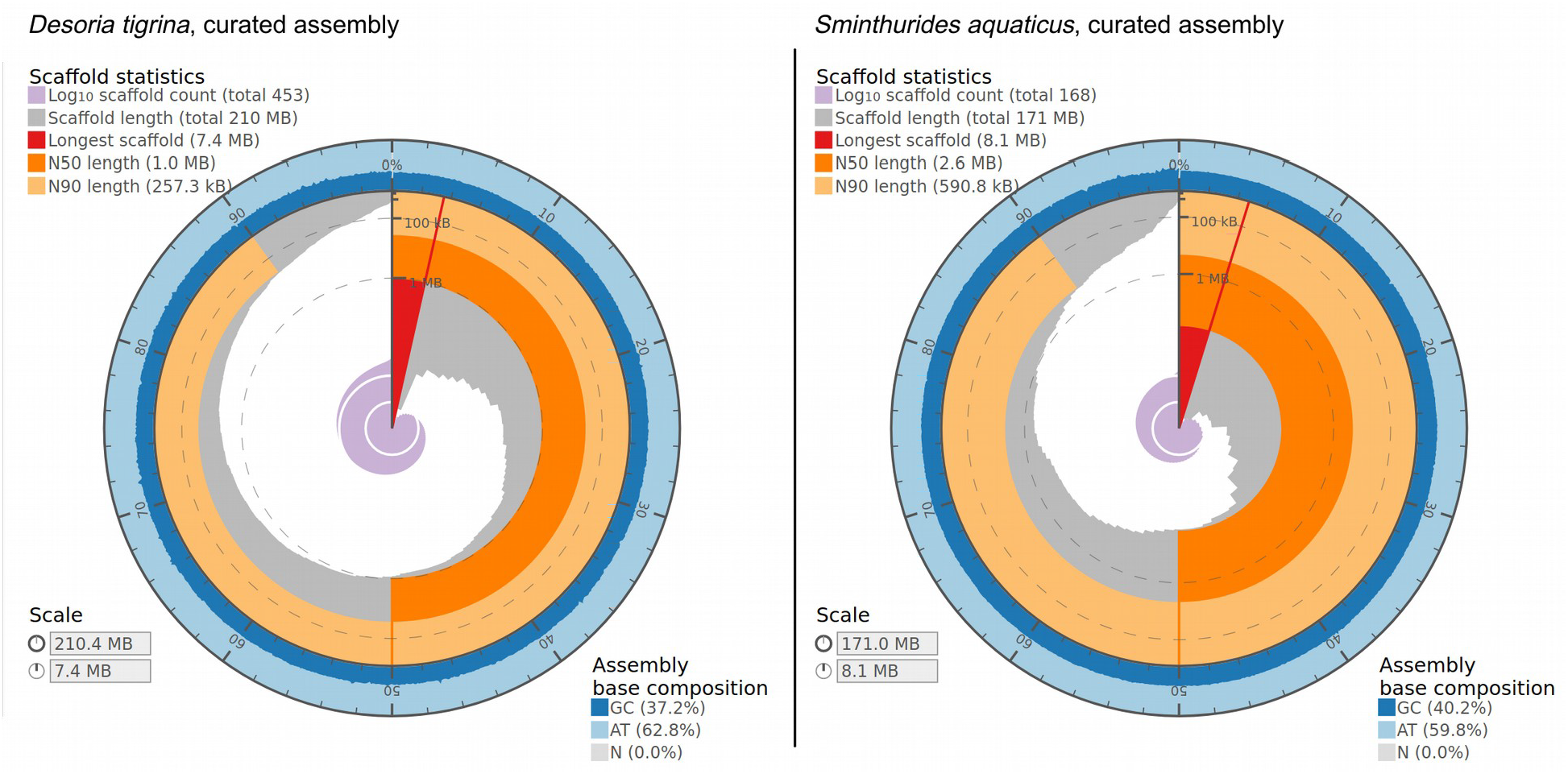
Curated assemblies visualization. Inner radius represents the length of the longest contig.

### Genome annotation

We predicted 17807 proteins in the genome of *S. aquaticus*, 11593 of which had homologs in other organisms, 6204 were labeled “hypothetical protein”. We predicted 21503 proteins in the genome of *D. tigrina*, 13576 of which had homologs in other organisms and 7927 were labeled “hypothetical protein”. We found four predicted genes in the *S. aquaticus* assembly which were associated with the metallo-beta-lactamase superfamily, and four predicted genes related to aminopenicillanic acid acyl-transferase. Nucleotide and protein searches did not reveal genes homologous of the isopenicillin N synthase-like genes neither in the *S. aquaticus*, nor in the *D. tigrina* genome sequenced with the UL protocol.

### Contamination

The SourMash assignment did not match contigs to a viral, bacterial or fungal genome for any of the two species. For *S. aquaticus*, the DIAMOND + MEGAN-LR assignment and the BlobToolKit pipeline (Megablast + Diamond) were not in complete agreement. The BlobToolKit generally provided more precise taxonomy: 10 contigs either assigned to Plantae species or unassigned by DIAMOND + MEGAN-LR were assigned to Collembola by BlobToolKit pipeline. 2 contigs unassigned by the BolbToolKit pipeline were assigned to Insecta species by DIAMOND + MEGAN-LR. Four small contigs (> 77084 bp) were either without assignment or assigned to Fungi or Streptophyta. A larger contig (212931 bp) was identified as a Cyanobacteria genome fragment. Those 14 contigs were excluded from downstream analysis. For *D. tigrina*, no evidence for contamination was returned by any approaches.

### Mitochondrial genomes and NUMTs

The complete set of 37 mitochondrial genes (13 proteins, 22 tRNA and 2 rRNA coding genes) found in Hexapoda were annotated for both species. The size of the mitochondrial genomes are *S. aquaticus:* 16.099 bp, *D. tigrina:* 15.139 bp. The larger size of the genome of *S. aquaticus* compared to *D. tigrina* reflects more intergenic space, including a peculiarly large one between the 16S rDNA and tRNA(Ile) (1302 bp vs 506 bp respectively). One large NUMT was detected in *S. aquaticus*, and four in *D. tigrina*. Number and length of NUMTs found in both species is reported in Figure 3.

**Figure 3.**
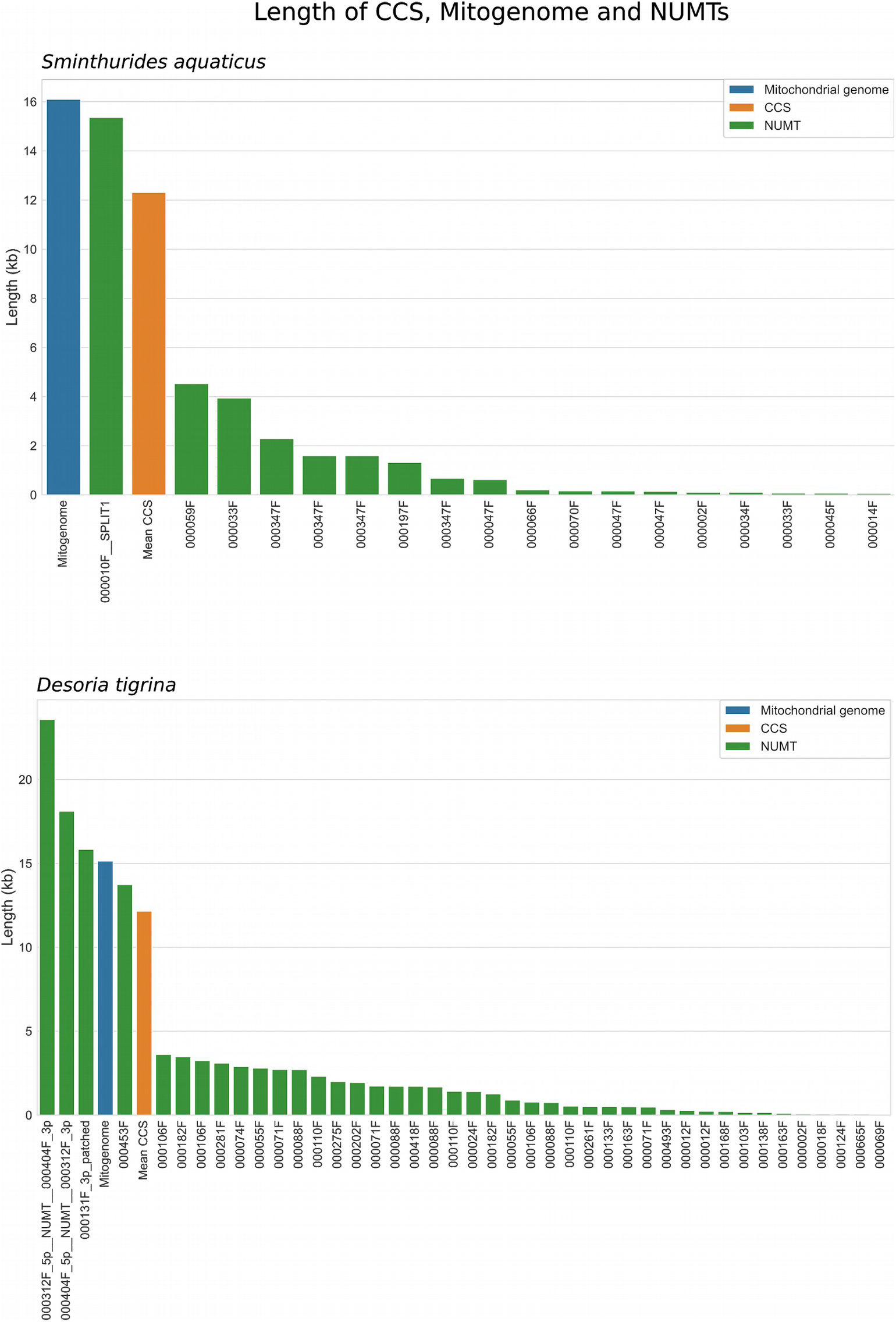
Length of the NUMTs (green) compared to the mitogenome (blue) and the mean CCS length (orange). The large NUMTs with length similar to the mitogenome creates assembly and polishing issues that were manually corrected. X labels are the name of the contigs where the NUMTs are observed.

### Phylogeny

We identified 932 single copy orthologs in the genome assemblies. After screening these for shared genes, we ended up with 597 protein sequences. Almost every divergence event has a bootstrap support of 100%, except the basal ones. Root attached on the branch linking Neelipleona to the rest of Collembola. The monophyly of Symphypleona, Poduromorpha and Entomobryomorpha is recovered. Symphypleona are sister-groups of Arthropleona (Poduromorpha, Entomobryomorpha). Within Entomobryomorpha, the represented super-family are monophyletic (two OTUs for each) and Tomoceroidea is sister-group of Isotomoidea + Entomobryoidea. Within Symphypleona, Arrhopalitidae are the sister-group of the rest. Sminthurididae (genus *Sminthurides*) are monophyletic and sister-group to Bourletiellidae and Sminthuridae). Within Poduromorpha, relationships are (Hypogastruridae + Neanuridae) + (Tullbergiidae + Onychiuridae). Tree is reported in Figure 4.

**Figure 4.**
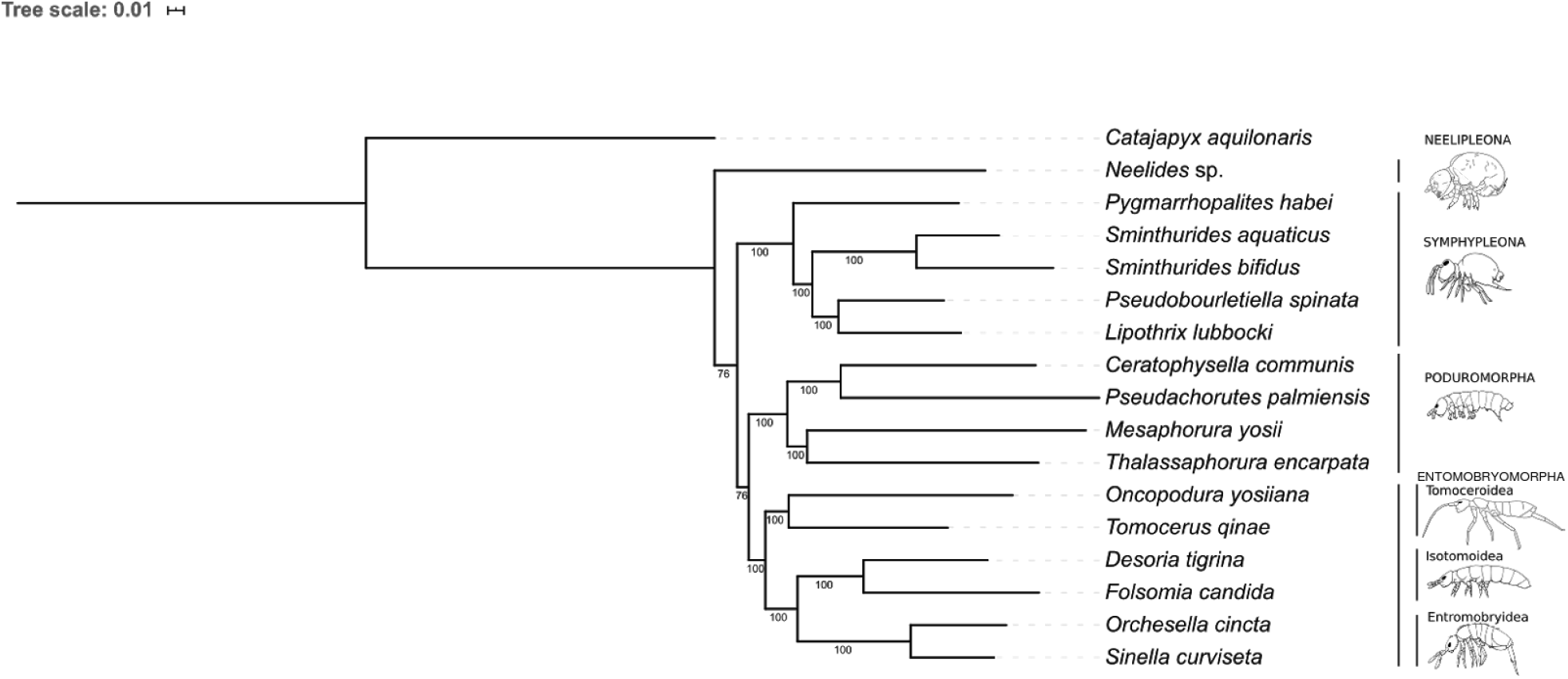
Phylogeny of Collembola based on the alignment of 597 protein sequences. Bootstrap supports are reported next to the nodes.

## DISCUSSION

We sequenced the reference genomes of two Collembola species, after we whole-genome-amplified high molecular weight DNA extracts from single specimens. Both species were field-collected and preserved in ethanol. The final assemblies were of high contiguity and completeness, on par with recent genomes sequenced on PacBio from single insects (Kingan et al., 2019b, 2019c), and also on par with the best reference genome for a Collembola so far, *Sinella curviseta*, which was DNA sequenced from 500 specimens maintained in culture (Zhang et al., 2019).

### Taxonomy

Field collected specimens in poorly known groups often present taxonomical issues, especially linked to cryptic diversity and complexity of species identification. The genus *Desoria* illustrates this complexity: within the *D. tigrina* group sensu Fjellberg 2007, *D. tigrina* and *D. grisea* (Lubbock, 1869) are two sibling species, described by pioneers of modern Collembola systematics. *Desoria grisea* was redescribed by Fjellberg (2007) from its type locality. It is distinguished from *D. tigrina* by consistent chaetotaxy characters on the labial palp. We examined several specimens from our collection spot and recognized all the characters of *D. tigrina*. Sequencing from a single specimen prevented the risk of sequencing a chimeric species.

*S. aquaticus* was originally described from France and has been recognized to be widely spread throughout the Holarctic region. We confirmed that our specimens are identical to the accepted descriptions of *S. aquaticus*. S. *aquaticus* was originally described by Bourlet in 1841 probably from the north of France. Bourlet did not make any reference to a type series, and to our knowledge did not preserve any specimens. The population we sampled in Paris is ideal to provide a new reference for this species: the population is abundant, settled, and easily accessible for further studies. The designation of a neotype accompanied with abundant material – specimens in slides; in ethanol and high quality genomic information – makes for a fine example of integrative taxonomy.

### Genomes

The higher level of heterozygosity in *D. tigrina* compared to *S. aquaticus* seems consistent with the expected level of isolation of the populations. *D. tigrina* invaded the compost that was set up one year before the collection. The species is very mobile, being rather large and equipped with a long furca, and gene flow must be active across the nearby surrounding fields and gardens. On the other hand, the sampled population of *S. aquaticus* seems rather isolated in a small area (artificial pond in a public garden). The L assembly of *S. aquaticus* (12 pooled females) is closest to estimated size from the CCS kmer analysis, and yields less duplicated BUSCOs, without any haplotypes purging step. This is possibly explained by more pronounced collapsing of the haplotypes during this assembly based on high-error rates CLR. The haplotig-purged assemblies size for both species are superior to the kmer estimated genome size. The estimated 152,357 Mbp of *S. aquaticus* seems to be a slight underestimate, given a significant drop in BUSCOs discovery observed in the <160 Mbp CLR assembly done with Falcon (table 2). The final assembly of 210.36 Mbp of *D. tigrina* is significantly above the estimated size (167.82 Mbp), and could be imperfectly haplotigs-purged as hinted by the recovery of 3.8% duplicated BUSCOs.

Standing alone, the ultra-low protocol can already provide high quality genomes on par with libraries produced without genome-wide amplification. Comparison and combination of the ultralow and low input data sets for *S. aquaticus* led to two observations: both performed equally in recovering BUSCOs and the ultra-low approach allowed to produce an assembly of higher contiguity. The reads of the low input protocol were relatively short and produced in CLR mode, which can account for the lowest performances. Our results show that the ultra-low protocol does not systemically miss large fractions of the genome and it can be used as a reasonable stand alone approach for single specimen sequencing of small metazoans. The presence of large NUMTs in the nuclear genome led to misassemblies because they can hardly be fully crossed by ~10-12 kb reads and because those regions are extremely similar between each other and to the mitochondrial DNA. This was observed in both species and such large NUMTs could be common in springtails genomes. Combination with long-range positional information could improve assembly by scaffolding and detection of misassemblies.

We could not identify beta-lactam antibiotic genes in the two annotated genomes. This is not surprising in the case of *S. aquaticus* which does not need to cope with such high microbial load. Indeed, three of four analysed species from the springtail order Symphypleona do not possess these genes (Suring et al., 2017). It is surprising however, that we did not find evidence of these genes in the genome if *D. tigrina*, which inhabits soils and likely needs to cope with high microbial load. Suring et al. (2017) identified beta-lactam gene clusters in four of six Entomobryomorpha species they investigated, including *F. candida*, which belongs to the same family as *D. tigrina* (Isotomidae). Beta-lactam gene clusters are sometimes not recovered from the genomes of hemiedaphic species (Suring et al., 2017), and *D. tigrina* also belongs to this group (Potapov, 2001). It is not yet clear if these genes are indeed missing from the genomes of *D. trigina*, or if their detection needs more thorough analyses.

### Phylogeny

Our two newly sequenced species find their expected placement: *S. aquaticus* next to *S. bifidus* (both representants of the genus *Sminthurides*, family Sminthurididae), and *D. tigrina* next to *F. candida* (both representants of the family Isotomidae). Our phylogeny has two points of disagreements with the phylogeny of Sun et al. (2020) from which the present work draws the species dataset. First is the relationships between the orders of Collembola: Neelipleona are found to be the sister-group of the other order of Collembola in both studies, but while we have Symphypleona sister-group of Poduromorpha + Entomobryomorpha (=Arthropleona), their study supports Poduromorpha sister-group of Entomobryomorpha + Symphypleona. Second is the monophyly of Tomoceroidea (recovered in our study, not in Sun et al. 2020). Regarding the Tomoceroidea, they receive high bootstrap support from our analysis, and the competing hypothesis presented in Sun et al. (2020) is not strongly supported.

The disagreement in basal relationships is not unexpected, as it occurs in nodes with poor support scores. We do not claim that our extended dataset is relevent in addressing this issue. Both histories of Collembola evolution is conflicting with the traditional hypotheses based on morphological evidences, and peculiarly the body segmentation: viewed as ancestral in Poduromorpha (all trunk segments distinct), modified in Entomobryomorpha (reduction of the first thoracic segment and sometimes fusion of abdominal segment), and further modified in Neelipleona and Symphypleona (more or less complete coalescence of thoracic and abdominal segmentation and acquisition of a globular appearance). The Symphypleona group was initially created including the genera of Neelipleona (Börner 1901). Massoud (1971, 1976) separated them by erecting the order Neelipleona, arguing that the globular morphology of both groups was acquired through a different process (abdomen reduced in Neelipleona, inflated in Symphypleona). While molecular-based phylogenies have since a long time confirmed the ancient separation between Neelipleona and Symphypleona *sensu* Massoud, one should be cautious in definitely rejecting a potential sistership between the two groups. Basal relationships of Collembola are poorly supported and sensitive to data sampling. Effects of long branch attraction and random root positioning are likely to occur, and have been briefly discussed in Schneider, Cruaud & D’Haese (2011). On the other hand, the advanced coalescence of body segments (regardless of thoracic / abdomen volume ratio) offers a trivial support for Symphypleona + Neelipleona monophyly. A rather large set of other “suspect” similarities between Symphypleona and Neelipleona exists, but are also trivially inconclusive because impossible to polarize (the gutter-like mucro) or because of parallel evolution (either convergent acquisition or multiple secondary loss): neosminthuroid chaetae, wax rods secretion, absence of postantennal organ, furcal posterior spines. It is worth noting that our tree disagrees on the traditional view on Collembola basal relationships only by the root position (see Schneider, Cruaud & D’Haese, 2011), and further work on Collembola phylogenomics should focus on improving the outgroup dataset.

### UL and L comparison

The comparison of genome completeness between the ultra-low input and low input protocol further suggests that the assemblies obtained from the ultra-low input pipeline are of the highest quality that could be expected. Unfortunately, the rather short reads yielded by our low input sequencing limits an in-depth comparison. Nonetheless, the almost identical BUSCO scores indicate that the genome-wide amplified ultra-low sequencing does not result in a biased representation of the genome, which is one of the main concerns of sequencing including a PCR step. Although PacBio sequencing remains more expensive than its short-read sequencing e.g. with Illumina, the more expensive long, but high fidelity (HiFi) reads provide a viable standalone solution to generate reference genomes. Our results demonstrate that long PacBio reads can now be used to sequence genomes of a much wider part of the Tree of Life, extending biodiversity genomics research to many small metazoans around the one millimeter size range.

Some highly interesting terrestrial taxonomic and functional groups in this regard are beetles, Diptera Hymenoptera, mites, and the soil mesofauna. Beetles were long considered the most biodiverse animal group (Hutchinson, 1959; Zhang et al., 2018), with amazing adaptations and tremendous diversity in the tropics. However, recent estimates suggest that both Diptera (Brown et al., 2018) and Hymenoptera (Forbes et al., 2018) might be considerably more diverse than beetles. This is due to a huge number of undescribed small species. Arachnids are also highly diverse, with potentially over 1 million species, most of which are mites (Stork, 2018). One of the largest reservoirs of terrestrial biodiversity is the soil mesofauna. Those animals, typically defined by sizes between 0.1—2 mm, represent many ancient terrestrial lineages: nematodes, potworms, mites, springtails, proturans, pauropods, insects, isopods, myriapods among others. Small metazoans are important contributors of ecosystem services (pollination, pests and parasites, biocontrol agents, etc. (Scholes, 2018). Genome sequencing is increasingly relevant for monitoring and managing these communities, from detecting underground invasions (Andújar et al., 2017) to chemical-free pest control. For example, genome information is essential to identify targets for RNA interference, an emerging species-specific alternative to chemicals (Vogel et al., 2019). Small metazoans are also important sources of novel natural products (Vilcinskas, 2014). Information on these products might be encoded in the genomes of the metazoans themselves (Faddeeva-Vakhrusheva et al., 2017; Drukewitz & von Reumont, 2019), or in the microbial symbionts which are co-sequenced with their hosts (Agamennone, 2019; Tobias, 2016). Genome-sequencing small metazoans will provide insights into the formation and maintenance of eukaryotic biodiversity and this will open up opportunities for natural resource management and bioprospecting.

## Acknowledgements

The present study is a result/product of the Centre for Translational Biodiversity Genomics (LOEWE-TBG) and was supported/funded through the programme “LOEWE – Landes-Offensive zur Entwicklung Wissenschaftlich-ökonomischer Exzellenz” of Hesse’s Ministry of Higher Education, Research, and the Arts. We thank the Genome Technology Center (RGTC) at Radboudumc for the use of the Sequencing Core Facility (Nijmegen, The Netherlands), which provided the PacBio SMRT sequencing service on the Sequel II platform. We highly appreciate the generous support by Pacific Bioscience with respect to the ultra-low amplification kit, library preparation kit as well as SMRT cells and sequencing chemistry during the course of the beta test. The Max-Planck Genome Center Cologne acknowledges the support from the Max-Planck Society. We give our warm thanks to Damian Baranski for supporting the lab work and DNA extractions in LOEWE-TBG Laboratory Center.

## DATA ACCESSIBILITY

The project is deposited in the EMBL-ENA database under accession number PRJEB39696 including: *S. aquaticus* CCS, UL raw reads and assembly and annotation under accessions numbers ERR4407379, ERZ1473260, ERZ1473263, CLR and L assembly under accession numbers ERR4405416, ERZ1473261, *D. tigrina* CCS, UL assembly and annotation under accession numbers ERR4407422, ERZ1473259, ERZ1473262. (Data remain confidential until publication in a peer-reviewed journal).

## AUTHOR CONTRIBUTIONS

C. Schneider conceived the ideas, C. Schneider, M. Bálint and B. Hüttel designed methodology; C. Schneider and C. D’Haese collected, identified and photographed the specimens; B. Hüttel performed the ultra-low input protocol and sequencing, C. Greve did the PacBio low input library preparation; C. Woehle and C. Schneider assembled and analysed the genomes; Magnus Wolf and Axel Janke contributed the phylogenomic analysis. C. Schneider and M. Bálint led the writing of the manuscript. All authors contributed to the manuscript and gave final approval for publication.

